# Structural conservation and diversity of PilZ-related domains

**DOI:** 10.1101/814665

**Authors:** Michael Y. Galperin, Shan-Ho Chou

## Abstract

The widespread bacterial second messenger cyclic diguanylate (c-di-GMP) regulates a variety of processes, including protein secretion, motility, cell development, and biofilm formation. C-di-GMP-dependent responses are often mediated by its binding to the cytoplasmic receptors that contain the PilZ domain. We present here comparative structural and sequence analysis of various PilZ-related domains and describe three principal types of them: (i) the canonical PilZ domain, whose structure includes a six-stranded beta-barrel and a C-terminal alpha-helix; (ii) an atypical PilZ domain that contains two extra alpha-helices and forms stable tetramers, and (iii) divergent PilZ-related domains, which include the eponymous PilZ protein and PilZN (YcgR_N) and PilZNR (YcgR_2) domains. We refine the second c-di-GMP binding motif of PilZ as [D/N]hSxxG and show that the hydrophobic residue *h* of this motif interacts with a cluster of conserved hydrophobic residues, helping maintain the PilZ domain fold. We describe several novel PilZN-type domains that are fused to the canonical PilZ domains in specific taxa, such as spirochetes, actinobacteria, cellulosolytic clostridia, deltaproteobacteria, and aquificae. We propose that the evolution of three major groups of PilZ domains included (i) fusion of *pilZ* with other genes, which produced Alg44, cellulose synthase, and other multidomain proteins; (ii) insertion of a ∼200-bp fragment, which resulted in the formation of tetramer-forming PilZ proteins, and (iii) tandem duplication of *pilZ* genes, which led to the formation of PilZ dimers and YcgR-like proteins.

**IMPORTANCE:** Cyclic di-GMP is a ubiquitous bacterial second messenger that regulates motility, biofilm formation and virulence of many bacterial pathogens. The PilZ domain is a widespread c-di-GMP receptor that binds c-di-GMP through its RxxxR and [D/N]xSxxG motifs; some PilZ domains lack these motifs and are unable to bind c-di-GMP. We used structural and sequence analysis to assess the diversity of PilZ-related domains and define their common features. We show that the hydrophobic residue *h* in the second motif is highly conserved; it may serve as a readout for c-di-GMP binding. We describe three principal classes of PilZ-related domains: canonical, tetramer-forming, and divergent PilZ domains, and propose the evolutionary pathways that led to the emergence of these PilZ types.

## INTRODUCTION

Cyclic bis(3’→5’) dimeric guanosine monophosphate (c-di-GMP) is a widespread bacterial second messenger that regulates a variety of processes, including DNA repair, transcription, mRNA decay, motility, protein and polysaccharide secretion, cell development, and biofilm formation (1-4). Cyclic di-GMP has a well-documented contribution to virulence of such important human pathogens as *Vibrio cholerae, Pseudomonas aeruginosa, Salmonella enterica, Borrelia (Borreliella) burgdorferi*, and *Yersinia pestis* and plant pathogens such as *Xanthomonas campestris* and *X. citri* (1, 3, 5-7). Cellular responses to c-di-GMP-dependent signaling are mediated by specific receptors, which include two types of riboswitches and a wide variety of c-di-GMP-binding proteins. The structures and ligand-binding properties of experimentally characterized c-di-GMP receptor proteins have been the subject of several reviews (8-10). In the past several years, novel c-di-GMP-binding proteins have been discovered, with some being widespread in bacteria, such as MshEN (11, 12), and some showing relatively narrow phylogenetic distribution (13, 14). There are reasons to believe that the current list is still incomplete (9).

The ∼110-aa PilZ domain (<u>PF07238</u> entry in the Pfam database, ref. 15), was the first c-di-GMP receptor to be identified (16), owing to its presence at the C-terminus of the cellulose synthase catalytic subunit BcsA from *Acetobacter xylinum* (current name, *Komagataeibacter xylinus*), the protein whose studies led to the original discovery of the c-di-GMP (17, 18). That work also described a number of other PilZ-containing domain architectures, including PilZ dimers, fusions of PilZ with the N-terminal PilZN (referred to as YcgR domain, <u>PF07317</u>, in Pfam) and PilZNR (YcgR_2, <u>PF12945</u>, in Pfam) domains, a fusion with HlyD family secretion protein in the alginate biosynthesis protein Alg44, as well as fusions of PilZ with REC, GGDEF, EAL, HD-GYP, chemoreceptor (MCPsignal), and CheC signaling domains (16).

Subsequent studies of various PilZ-containing proteins confirmed the ability of this domain to bind c-di-GMP and control a variety of c-di-GMP-regulated functions (19-28). It has become clear that different PilZ-containing proteins bind c-di-GMP with a wide range of affinities, which contributes to the diversity of c-di-GMP signaling mechanisms (29). Curiously, the eponymous PilZ protein of *P. aeruginosa* (genomic locus tag PA2960) does not bind c-di-GMP (21, 30). Instead, it appears to regulate formation of type IV pili through protein-protein interactions. In *Xanthomonas* spp., PilZ protein interacts with FimX (31, 32), while in *P. aeruginosa*, FimX does not do so and the PilZ-interacting partner remains to be characterized (33).

Over the years, structures of several PilZ domains and PilZ-containing proteins have been solved (22, 24, 25, 34-40), providing valuable insight into the mechanisms of c-di-GMP binding and the roles of the conserved RxxxR and [D/N]xSxxG c-di-GMP-binding motifs (reviewed in ref. 9 and 10). Here, we present a comparative analysis of PilZ sequences and structures, which allowed identification of additional conserved elements in PilZ domains and delineation of three major classes of PilZ-related domains. Based on this analysis, we propose a general scheme of PilZ evolution that involves three major evolutionary routes leading to (i) multidomain fusion proteins with canonical PilZ, (ii) tetrameric PilZ proteins, and (iii) two-domain proteins consisting of diverse PilZ-related N-terminal domains fused to the C-terminal canonical PilZ. Evolution along these routes is sometimes accompanied by the loss of c-di-GMP binding motifs.

## RESULTS

### Structural conservation of canonical PilZ domains

The typical PilZ domain core consists of six β-strands followed by a single long α-helix. These β-strands are arranged in two antiparallel β-sheets that form a β-barrel. In the past 12 years, more than 30 structures of various PilZ domains have been solved and deposited in the Protein DataBank (PDB), see Table S1. These include high-resolution crystal structures of the stand-alone PilZ domains, as well as PilZ domains from the bacterial cellulose synthase (BcsA and AcsAB), alginate production protein Alg44, flagellar brake proteins YcgR, VCA0042, and MotI, and the transcriptional regulator MrkH (22, 25-27, 34, 35, 38, 39, 41-43). As noted in the respective reports, most of these domains are very similar in sequence and structure. Indeed, comparison of representative PilZ structures using the Dali server (44) revealed that, despite their relatively low sequence identity (≤ 20%), these domains readily align over their entire lengths, with root mean standard deviation (RMSD) of the Cα traces within 3.2 Å (Table 1). Comparison of the same structures using the VAST tool, which computes an alignment of secondary structure elements, α-helices and β-strands (45), brought similar results, with slightly shorter aligned regions but a bit higher sequence identity and even lower RMSD values (≤ 2.5 Å, see Table S2).

**Table 1.** Structural similarity of canonical, tetrameric, and divergent PilZ domains.

| Organism, protein name<br>(genomic locus tag)<sup>a</sup> | PDB entry,<br>resolution | Alignment length,<br>% identity, RMSD<sup>b</sup> | Reference<sup>c</sup> |
| --- | --- | --- | --- |
| **Canonical** |  |  |  |
| *Pseudomonas aeruginosa* PilZ<br>or MapZ (PA4608) | 5XLY_B, 1.76 Å | 109 aa, 100%, 0 Å | 40 |
|  | 2L74_A, NMR | 121 aa, 100%, 2.1 Å | 36 |
| *Legionella pneumophila* PilZ<br>(Lpg0364) | 4Q63_A, 1.95 Å | 88 aa, 11%, 2.7 Å | N/A, MCSG |
| *Pseudomonas aeruginosa* PilZ<br>from Alg44 (PA3542) | 4RT1_B, 1.7 Å | 101 aa, 17%, 2.6 Å | 37 |
| *Komagataeibacter xylinus* PilZ<br>from AcsAB | 4I86_B, 2.10 Å | 92 aa, 20%, 2.6 Å | 43 |
| *Rhodobacter sphaeroides* BcsA<br>(RSP_0333, aa 578-691) | 5EJZ_A, 2.95 Å | 98 aa, 12%, 2.6 Å | 25 |
| *Escherichia coli* YcgR<br>(b1194, aa 111-244), | 5Y6G_A, 2.30 Å | 101 aa, 17%, 2.7 Å | 49 |
| *Pseudomonas putida* YcgR or<br>FlgZ (PP_4397, aa 121-238) | 3KYF_A, 2.10 Å | 101 aa, 12%, 2.5 Å | 41 |
| *Vibrio cholerae* PlzD<br>(VCA0042, aa 134-247) | 1YLN_A, 2.20 Å | 99 aa, 14%, 3.7 Å | N/A,<br>MCSG;<br>22 |
|  | 2RDE_A, 1.92 Å | 99 aa, 14%, 3.1 Å |  |
| *Bacillus subtilis* MotI or DgrA<br>(BSU22910, aa 97-208) | 5VX6_A, 3.2 Å | 104 aa, 16%, 3.2 Å | 39 |
| *Klebsiella pneumoniae* MrkH<br>(KP13_02245, aa 107-234) | 5KEC_D, 1.95 Å | 104 aa, 11%, 3.1 Å | 38 |
| **Tetrameric (tPilZ)** |  |  |  |
| *Xanthomonas campestris*<br>Xcc0612 (XccB100_2234) | 3RQA_A, 2.1 Å | 93 aa, 12%, 2.9 Å | 46 |
| **Divergent (xPilZ)** |  |  |  |
| *Xanthomonas campestris* or *X.*<br>*citri* PilZ (Xcc1024, Xac1133) | 3CNR_B, 1.90 Å | 72 aa, 13%, 2.9 Å | 34, 35 |
|  | 3DSG_B, 2.09 Å |  |  |
| *Geobacter sulfurreducens*<br>GSU3033 (aa 1-102) | 2L1T_A (NMR) | 85 aa, 5%, 3.1 Å | N/A, JCSG |
| *Escherichia coli* YcgR_N<br>(b1194, aa 1-110) | 5Y6H_A, 1.77 Å | 80 aa, 4%, 2.9 Å | 49 |
| *Vibrio cholerae* PlzD<br>(VCA0042, aa 24-130) | 2RDE_A, 1.92 Å | 75 aa, 9%, 2.2 Å | 22 |
| *Klebsiella pneumoniae* MrkH<br>(KP13_02245, aa 1-106) | 5KEC_D, 1.95 Å | 77 aa, 10%, 2.9 Å | 38 |

- <sup>a</sup> – For multidomain structures, the amino acid (aa) boundaries of the respective PilZ domains are shown in parentheses. - <sup>b</sup> – The overlap length, percent identity, and RMSD values were taken from DALI and/or VAST (44, 45) alignments of the respective structures against the stand-alone PilZ domain in the *Pseudomonas aeruginosa* protein MapZ (PDB: 5XLY\_B). - <sup>c</sup> – N/A, unpublished. MCSG, Midwest Center for Structural Genomics; JCSG, Joint Center for Structural Genomics.

Structural superposition of PilZ domains from *K. xylinus* AcsAB, *P. aeruginosa* Alg44 and C-terminal domains of *E. coli* YcgR, *V. cholerae* PlzD, and *K. pneumoniae* MrkH against the high-resolution structure of the PilZ domain protein MapZ (PA4608) of *P. aeruginosa* (24, 26, 40) is shown in Fig. 1, while pairwise superpositions of the domains listed in Table 1 are shown in Fig. S1. Structure-guided sequence alignment of these domains is presented in Fig. S2 and, in an abbreviated form, in Fig. 2. These comparisons demonstrate that, despite some differences in the lengths of β-strands and occasional split of a long β-strand into two, PilZ domains of the BcsA/AcsAB, YcgR, PlzD, MrkH, and Alg44 proteins adopt essentially the same core structure as stand-alone PilZ domains. Accordingly, they can all be considered “canonical” PilZ domains.

**Figure 1.**
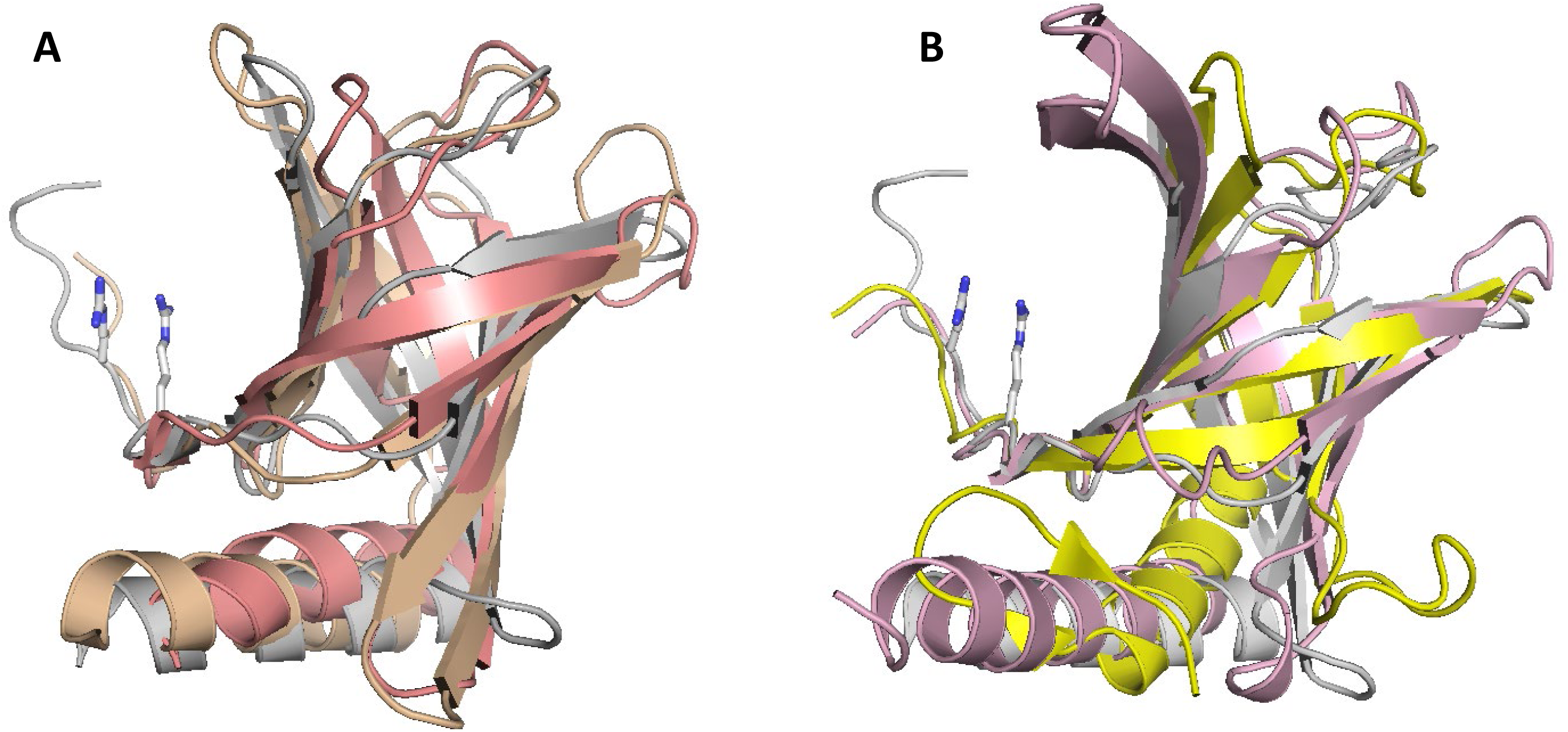
Structural superposition of canonical PilZ domains. **A.** Stand-alone PilZ domain protein MapZ (PA4608) from *P. aeruginosa* (PDB entry 5XLY_B, residues 4-107), shown in grey, superposed against PilZ domains from *K. xylinus* cellulose synthase (PDB: <u>4I86</u>, shown in salmon) and from *P. aeruginosa* Alg44 (PDB: <u>4RT1</u>, shown in wheat color). **B.** MapZ structure (in grey) superposed against C-terminal PilZ domains of *E. coli* YcgR (PDB: <u>5Y6G</u>, shown in light pink) and *V. cholerae* PlzD (PDB: <u>2RDE</u>, residues 134-237, shown in yellow). The Arg residues of the MapZ RxxxR motif are in stick representation.

**Figure 2.**
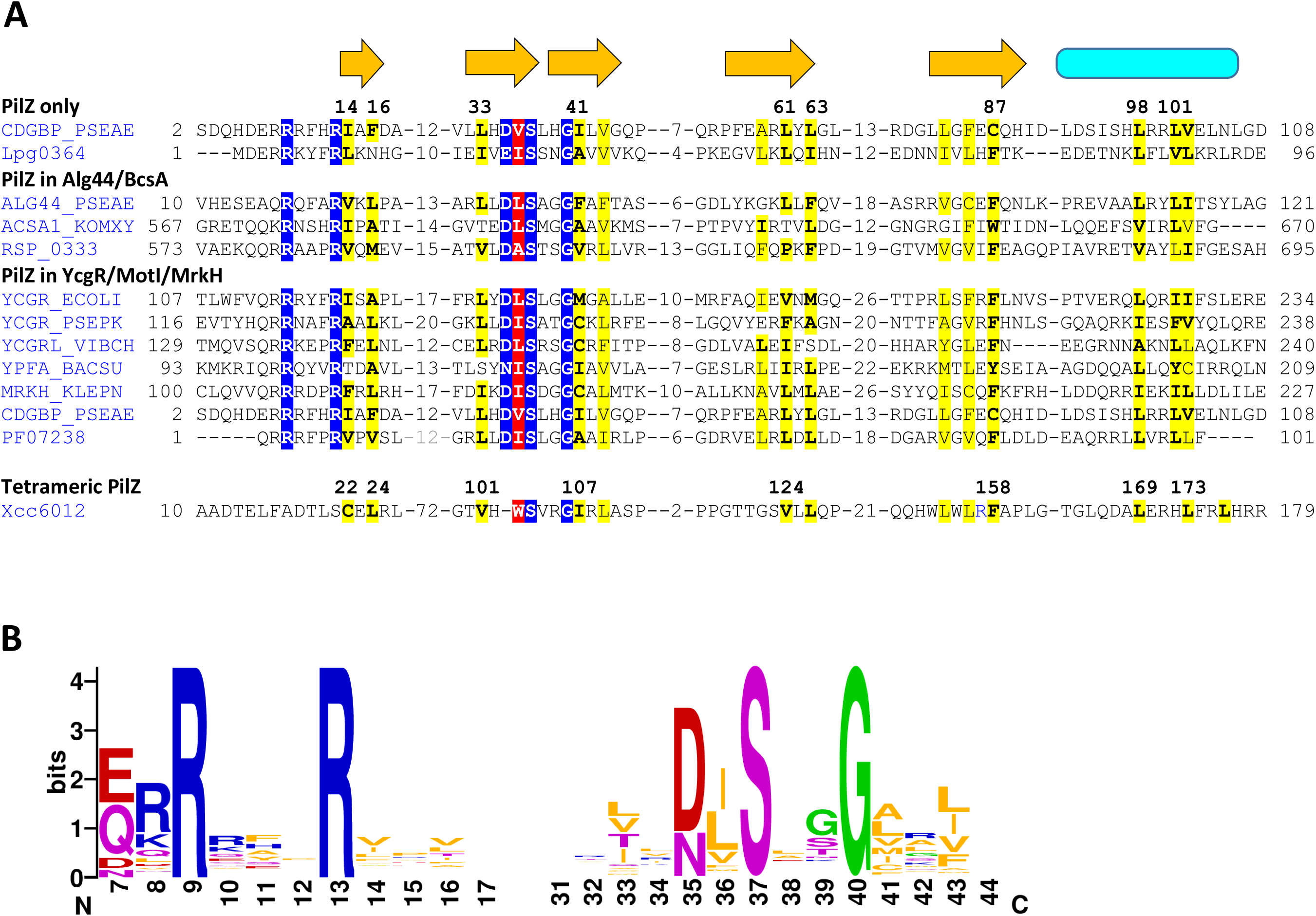
Sequence conservation of canonical PilZ domains. **A.** Structure-based sequence alignment of canonical PilZ domains. The residues of the c-di-GMP-binding motifs RxxxR and [D/N]xSxxG are in white on blue background. The proteins are listed under their UniProt names or genomic locus tags. The secondary structure elements (β-strands, arrows, and the α-helix, cylinder) are from the *P. aeruginosa* MapZ structure (PDB: 5XLY_B). Conserved hydrophobic residues are shaded yellow, those forming the hydrophobic pocket are shown in bold and labeled by their numbers in the MapZ structure. The newly identified conserved *h* residue of the [D/N]hSxxG motif is in white on red background. For comparison, the sequences of the PilZ domain entry in Pfam (PF07238) and the tetramer-forming tPilZ domain (PDB: 3RQA) are shown at the bottom. The complete alignment is presented in Fig. S3. **B.** Sequence logos of the c-di-GMP-binding motifs.

Another variation on the theme is the PilZ-PilZ domain architecture that represents tandem duplication of the canonical PilZ domain. In these proteins, exemplified by the previously described (16) Bll4394 and Blr5568 proteins from *Bradyrhizobium japonicum* (current name, *B. diazoefficiens*), both PilZ domains keep their C-terminal α-helices and often retain intact c-di-GMP-binding motifs. The current version of Pfam (15) lists more than 240 such PilZ-PilZ proteins, primarily from alphaproteobacteria but also some from gamma- and delta-proteobacteria, planctomycetes, and candidate division NC10. While analyzing PilZ fusion proteins (see below), we were able to substantially expand this list, with additional members again found primarily in alpha-proteobacteria. The largest such family includes *Sinorhizobium meliloti* protein SMc00999 and more than a thousand of closely related proteins. As discussed below, these proteins are important from the evolutionary point of view, as tandem duplications of canonical PilZ domain, followed by diversification of the N-terminal domain, give rise to a variety of YcgR-like two-domain proteins.

### Conservation of hydrophobic residues, the [D/N]hSxxG motif

In addition to the well-known RxxxR and [D/N]xSxxG motifs, the original description of the PilZ domain (16) showed conservation of two hydrophobic residues. One was located at the end of the second β-strand between the D/N and S residues of the second c-di-GMP-binding motif and the other (mostly Phe) was near the end of the last β-strand (Leu/Ile/Val/Phe-32 and Phe/Trp/Tyr-90, respectively, in Figure 1 of ref. 16). Yet, the roles of these hydrophobic residues have not been investigated. Now, using structure-based sequence alignment of these domains, we were able to confirm strict conservation of these and several other hydrophobic residues (Fig. 2A). Therefore, the second c-di-GMP-binding motif of PilZ could be redefined as [D/N]hSxxG, where *h* stands for a hydrophobic residue, which is primarily Ile, Leu, or Val at this position (Fig. 2B). This observation prompted us to check the positions of this *h* residue in various PilZ structures.

As shown in Fig. 3, the [I/L/V] hydrophobic residue, while located within the c-di-GMP-binding motif, does not interact with c-di-GMP. Instead, its side chain is oriented towards the other side of the β-strand and is located within a hydrophobic pocket formed by at least 7 other hydrophobic residues. Sequence alignment (Fig. 2A) shows that the positions of these residues were well conserved among various PilZ domains, although the residues themselves varied. The most conserved of them was the previously noted Phe residue (replaced by Cys-87 in *P. aeruginosa* MapZ), located near the end of the last β-strand of PilZ (Fig. 2A). Comparison of the PilZ structures showed that this Phe residue directly interacts with *h* residue of the [D/N]hSxxG motif (Val-36 in *P. aeruginosa* MapZ) and apparently limits the mobility of the Gly residue of this motif. Another pocket-forming hydrophobic residue is located immediately after the Gly of the [D/N]hSxxG motif (Fig. 2). However, at that position, aliphatic residues (Ile, Leu, Val, Ala) were found along with Met and Cys (Fig. 2), which is why we refrained from including this second *h* into the definition of the c-di-GMP binding motif as [D/N]hSxxGh.

**Figure 3.**
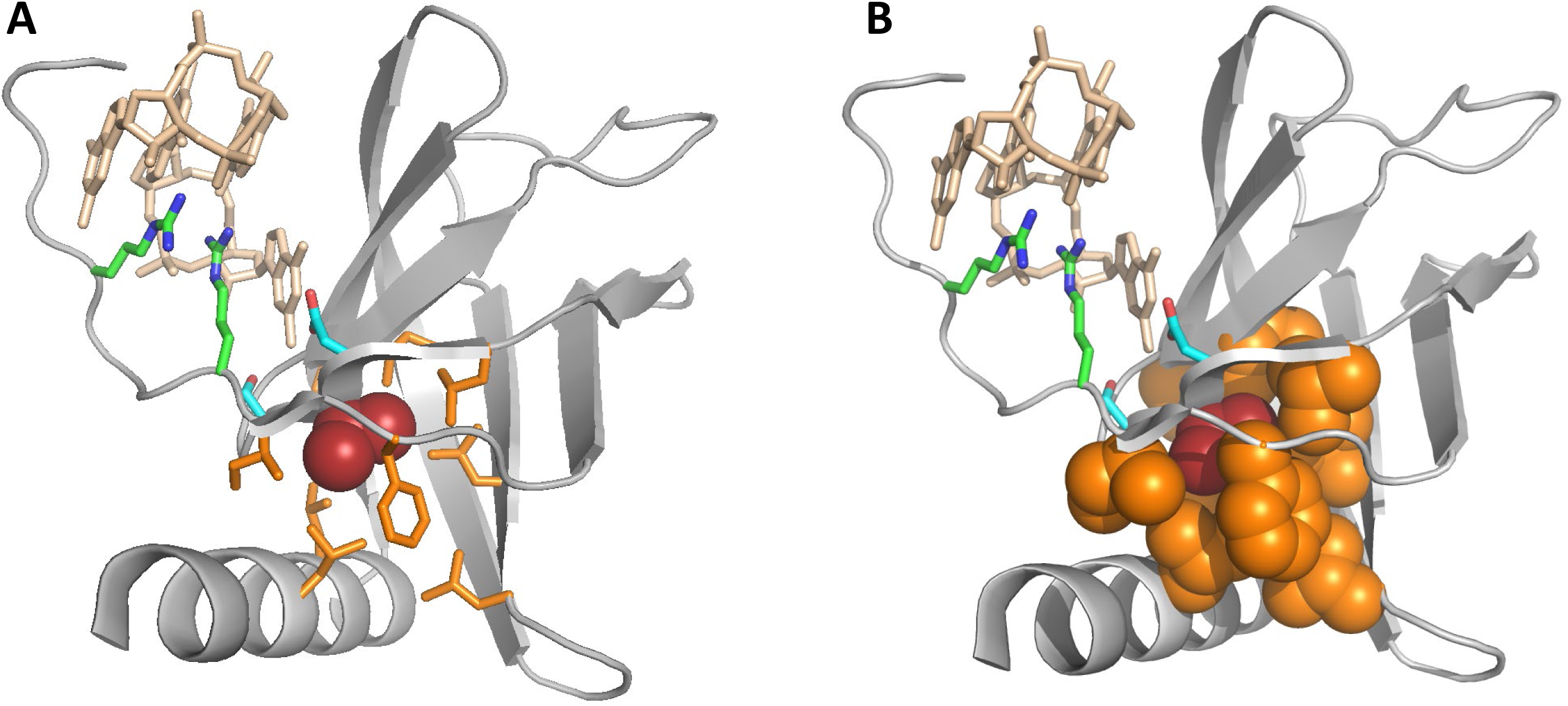
The hydrophobic pocket surrounding the *h* residue of the [D/N]hSxxG motif. The structure of the *P. aeruginosa* MapZ (PDB: 5XLY_B) is in light grey; the c-di-GMP dimer is in wheat color, and the c-di-GMP-binding Arg9 and Arg13, Asp35 and Ser37 residues are shown in stick representation with C atoms, respectively, in green and blue. The *h* residue of the [D/N]hSxxG motif, Val36, is dark red. The Ile14, Phe16, Leu33, Ile41, Leu61, Leu63, Cys87, Leu98, and Leu101 residues (highlighted in Fig. 2), are in orange and shown in stick (A) or filled space (B) representation.

As seen in Fig. 2, two more residues that form the hydrophobic pocket are located after the RxxxR motif in the RxxxRhxh pattern. However, these two positions can be occupied by various uncharged (or even charged) residues, so we have chosen to leave the RxxxR motif as it is.

### Tetramer-forming PilZ domains

The Xcc0612 protein from *X. campestris* (GenBank accession <u>ALE68703</u>, PDB: <u>3RQA</u>) contains an “atypical” PilZ domain (PilZ_2, Pfam domain <u>PF16823</u>, referred to hereafter as tPilZ) that displays essentially the same structure as canonical PilZ (RSMD = 2.2 - 2.9 Å), with the exception of a short α-helix at the N-terminus and an ∼60-aa insert that forms two additional α-helices, α2 and α3 (46), see Fig. 4A. In the Xcc6012 structure, α3 helices from four PilZ protomers interact with each other via strong hydrophobic hepta-repeats, leading to the formation of a very stable tetramer, which is strengthened further by intermolecular bridges between Lys-77 in the middle of the ^75^DAKLD^79^ motif with Asp-75 and Asp-79 of the adjacent monomer (46). Previously, tPilZ domains were thought to be limited to xanthomonads. We have now detected the respective genes in the genomes of a wide variety of gamma-proteobacteria, as well as in certain members of other subdivisions of *Proteobacteria* and of the phyla *Nitrospirae, Nitrospinae/Tectomicrobia, Deferribacteres*, and *Thermodesulfobacteria* (Fig. S3, Table S3), with more than 3,000 proteins in the current protein databases. All these proteins have similarly-sized inserts that form two predicted α-helices but display certain variation of the domain-interlocking motif, which was generalized to [DENH]x[KR]h[DEN], where *h* again indicates a hydrophobic amino acid residue. Further, the tPilZ domain of Xcc0612 has disrupted RxxxR and [D/N]hSxxG motifs and is unable to bind c-di-GMP (46). By contrast, tPilZ domains from many other bacteria have these motifs intact, suggesting that they could still bind c-di-GMP. Indeed, *P. aeruginosa* protein PA2989 (which has a typical RxxxR motif but the [D/N] residue of the second motif is replaced by Ser, Fig. S3), has been experimentally shown to bind c-di-GMP (47, 48). In contrast, in PA2989 homologs from some other *Pseudomonas* species, these motifs are disrupted (Fig. S3), indicating the likely loss of c-di-GMP-binding within a single bacterial genus. Intact motifs are also present in the tPilZ protein Lpg1926 from the important pathogen *Legionella pneumophila* (Fig. S3), suggesting that it, too, has retained the ability to bind c-di-GMP.

**Figure 4.**
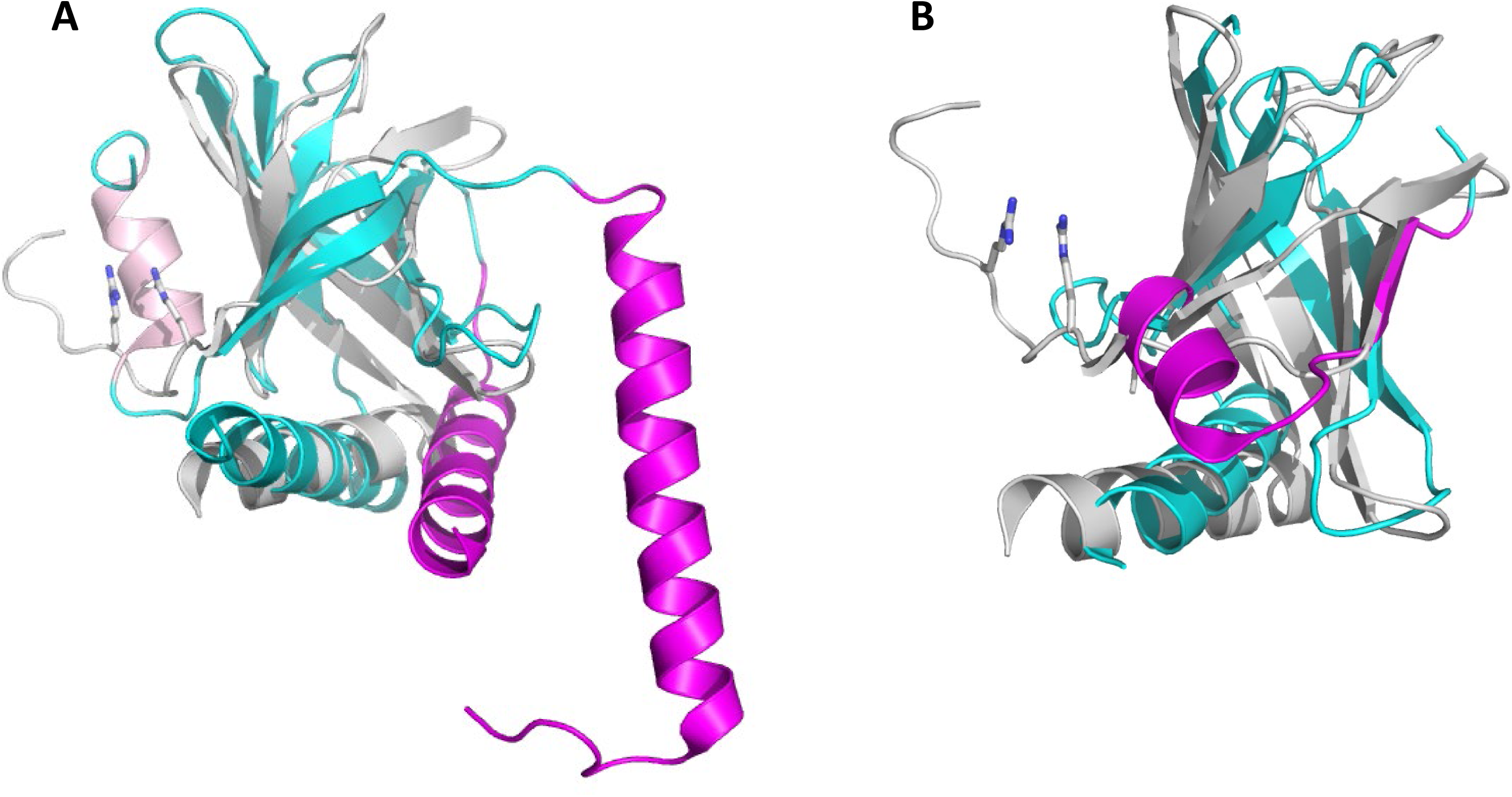
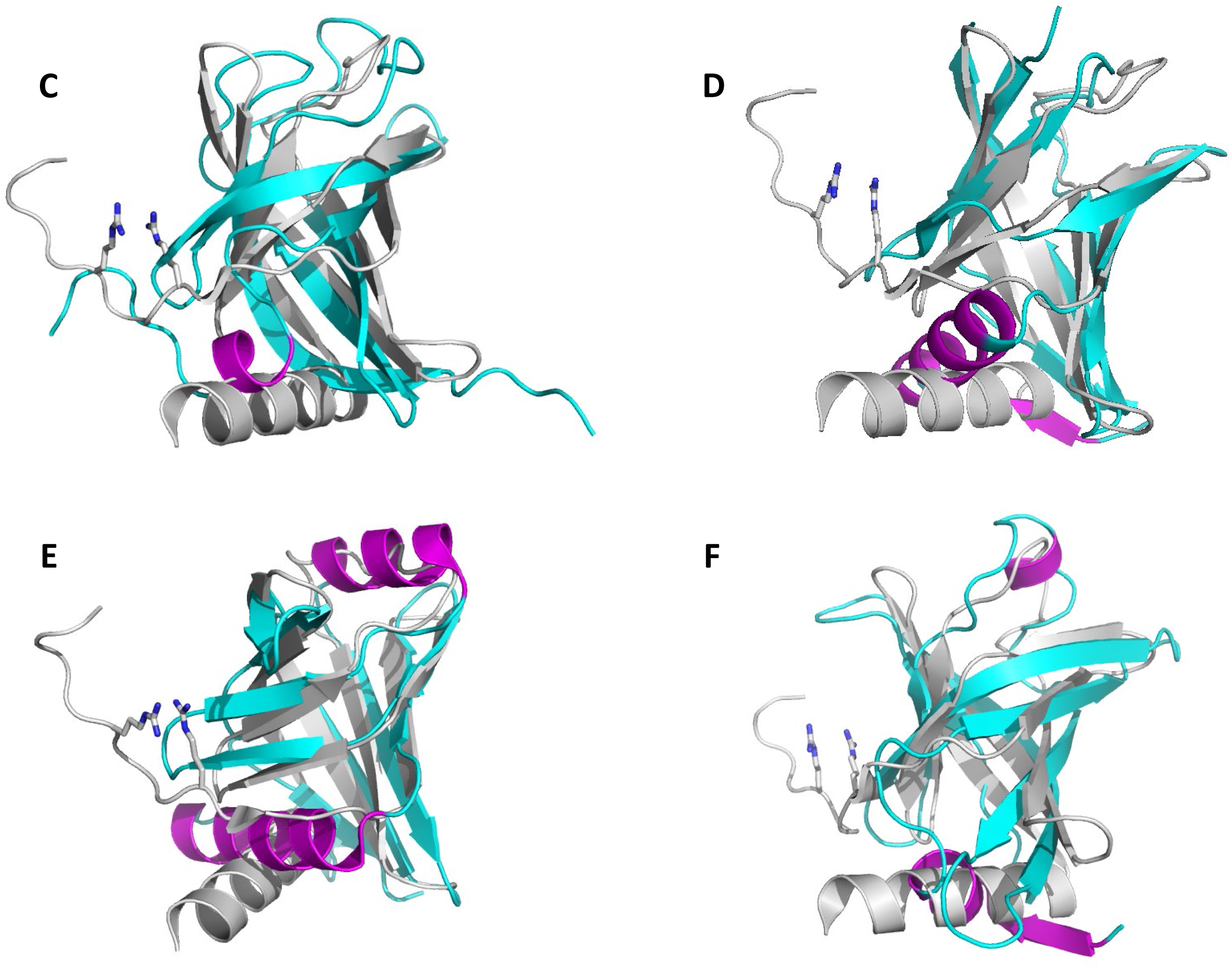
Structural similarity of PilZ-related domains. Canonical PilZ domain protein MapZ (PA4608) from *P. aeruginosa* (PDB entry 5XLY_B, residues 4-107) superposed against the tetramer-forming PilZ domain from *X. campestris* (PDB: 3RQA, panel A); the stand-alone c-di-GMP-nonbinding PilZ protein from *X. citri* (Xac1133, PDB: 3CNR, panel B); the DUF5634_N domain from *G. sulfurreducens* protein GSU3033 (PDB: 2L1T, panel C), and the N-terminal domains domains from *E. coli* YcgR (PDB: <u>5Y6H</u>, panel D), *V. cholerae* PlzD (YcgR_2, PDB: <u>2RDE</u>, residues 24-130, panel E), and *K. pneumoniae* MrkH (PDB: <u>5KEC</u>, residues 1-106, panel F). In all panels, the reference PilZ structure 5XLY is in light grey, aligned structural elements from other proteins are in cyan, structural elements that deviate from the canonical PilZ structure are in magenta. The Arg residues of the MapZ RxxxR motif are in stick representation.

Even when the [D/N]hSxxG motif is not conserved, tPilZ domains retain a hydrophobic residue (typically I/L/V but occasionally W or F) in the position of *h* and the constellation of hydrophobic residues to form a pocket around it (Fig. 2 and S3). One of these residues immediately precedes the (former) [D/N]hSxxG motif, whereas in canonical PilZ domains it is separated from this motif by an additional residue (cf. Fig. S2 and S3). tPilZ domains also retain the same amphipathic C-terminal α-helix that provides two hydrophobic residues to this pocket. This α-helix typically contains a patch of Arg, Lys, and His residues and carries a strong positive charge (Fig. S3).

In contrast to the canonical PilZ, tPilZ domains are almost never seen forming fusions to other domains. Besides, these domains always appear to be encoded in a single copy per genome, irrespective of the total number of other PilZ domain proteins encoded in that genome (Table S3). The reasons for this behavior remain to be investigated.

### Divergent PilZ-related domains

While the canonical and tetramer-forming PilZ domains share the same core structure of a six-stranded β-barrel followed by an α-helix, members of the third group, referred to here as “divergent” PilZ-related domains, exhibit various deviations from that pattern.

One of such deviations is seen in the stand-alone PilZ domain from two nearly identical proteins, Xc1028 from *X. campestris* and Xac1133 from *X. citri* (34, 35). These proteins lack the c-di-GMP-binding motifs and participate in the regulation of type IV pili formation through their interaction with the EAL domain of the FimX protein and the pilus motor protein PilB (31, 32, 34). The loss of the N-terminal loop with its RxxxR motif is accompanied by the inversion of the direction of the N-terminal β-strand and conversion of the second β-strand into an α-helix (34, 35). Accordingly, the five remaining β-strands do not form a complete barrel and the alignment of this structure against canonical PilZ is much shorter (72 aa residues) with an RMSD of 2.9 Å (Table 1, Fig. 4B). The sequence of the eponymous PilZ protein (PA2960) from *P. aeruginosa* (30, 35) is very similar (67% identity) to those of Xc1028 and Xac1133, indicating that it, too, belongs to the group of “divergent” PilZ domains (abbreviated hereafter as xPilZ).

The greatest variety of PilZ-related domains is found in a vast superfamily of two-domain proteins that combine various xPilZ domains at their N-termini with canonical PilZ domains at their C-termini. These proteins are generally similar to the previously described (16) alphaproteobacterial proteins with tandemly duplicated PilZ domains, such as *B. diazoefficiens* Bll4394 and Blr5568, but show various structural changes in their N-terminal PilZ-like domains. Based on the similar lengths and sequence similarity to the experimentally characterized flagellar brake proteins YcgR and MotI (22, 39, 41, 49), these proteins are often referred to as YcgR or YcgR-like. However, sequence similarity between these proteins is mostly limited to their common C-terminal PilZ domains. In contrast, their N-terminal domains exhibit various deviations from the PilZ core structure, as seen, for example, in the four structurally characterized domains: PilZN (YcgR, <u>PF07317</u>), PilZNR (YcgR_2, <u>PF12945</u>), DUF5634_N (<u>PF18672</u>) and the N-terminal domain of the transcriptional regulator MrkH, see Fig. 4, panels C-F. While all these domains have a β-barrel fold like the one in canonical PilZ, they all lack the C-terminal α-helix. They also show additional deviations from the PilZ-PilZ consensus of Bll4394 and Blr5568 proteins.

The least changes are seen in the GSU3033 protein of *Geobacter sulfurreducens* (GenBank: <u>AAR36425</u>), which contains a canonical PilZ domain at its C-terminus. The N-terminal domain of GSU3033 has been characterized by NMR (PDB: <u>2L1T</u>) and, along with related deltaproteobacterial domains, placed in a separate Pfam family DUF5634_N (<u>PF18672</u>). This domain, referred to PilZN1 in Table 2, has the same N-terminal long loop followed by a six-stranded β-barrel as the canonical PilZ domain but lacks the C-terminal α-helix and both c-d-GMP-binding motifs; in addition, the N-terminal loop forms a short (3-aa) α-helix (Fig. 4C).

**Table 2.** Non-canonical PilZ-related domains.

| Domain name,<br>Pfam entry<sup>a</sup> | Representative example |  |  | Phylogenetic<br>distribution |
| --- | --- | --- | --- | --- |
|  | Organism, locus tag<sup>b</sup> | GenBank<br>entry | Bound<br>aries |  |
| tPilZ (PilZ_2),<br>PF16823 | *V. cholerae* VC1885,<br>*P. aeruginosa* PA2989,<br>*T. indicus* Thein_0310 | AAF95033,<br>AAG06377<br>AEH44194 | 1-190,<br>1-230,<br>1-190 | Proteobacteria,<br>Nitrospirae, Thermo-<br>desulfobacteria |
| PilZN (YcgR),<br>PF07317 | *E. coli* YcgR,<br>*P. putida* PP_4397 | AAC74278,<br>AAN69975 | 1-110,<br>1-115 | Beta-, gamma-proteobacteria |
| PilZNR (YcgR_2),<br>PF12945 | *V. cholerae* VCA0042,<br>*B. subtilis* BSU22910 | AAF95956,<br>CAB14207 | 1-130,<br>1-90 | Proteobacteria,<br>Firmicutes, other |
| PilZN1, PF18672<br>(DUF5634_N) | *Geobacter sulfurreducens*<br>GSU3033 | AAR36425 | 1-105 | Desulfuromonadales |
| PilZN2 | *K. pneumoniae* MrkH<br>(KP13_02245) | AEO27488 | 1-105 | Enterobacteriales |
| PilZN3 | *B. burgdorferi* BB0733,<br>*T. denticola* TDE0214 | AAC67086,<br>AAS10711 | 1-140,<br>1-140 | Spirochetes |
| PilZN4 | *T. denticola* TDE1318<br>*A. aeolicus* Aq_1917 | AAS11835,<br>AAC07718 | 1-210,<br>1-220 | Spirochetes, Aquificae,<br>Fibrobacteres |
| PilZN5 | *G. obscurus* Gobs_0077,<br>Gobs_0078 | ADB72887 | 1-95 | Actinobacteria |
| PilZN6 | *Desulfarculus baarsii*<br>Deba_0512, Deba_3043 | ADK83884,<br>ADK86396 | 1-140,<br>1-140 | Deltaproteobacteria |
| PilZN7 | *H. thermocellum*<br>Cthe_0108, Cthe_0868 | ABN51349,<br>ABN52102 | 1-90,<br>1-100 | Hungateiclostridiaceae |

<sup>a</sup> – PilZN1-PilZN7 are tentative domain names, expected to be replaced as soon as these domains are experimentally characterized.
<sup>a</sup> – For complete names of the organisms, see the text or the respective GenBank entries.

The structure of MrkH, a transcriptional regulator of type 3 fimbriae production in *Klebsiella pneumoniae* (50), reveals yet another N-terminal six-stranded β-barrel domain (hereafter, PilZN2 domain) fused to a canonical PilZ domain (27, 38). In PilZN2, the C-terminal α-helix is lost, and the N-terminal loop is replaced by a short β-strand, followed by a 13-aa α-helix (Fig. 4D).

Like PilZN2, the PilZN (YcgR) domain structures from *E. coli* and *Pseudomonas putida* deviate from the canonical PilZ structure in the absence of the C-terminal α-helix and presence of a 13-aa α-helix after the β1 strand. In addition, they contain an 8-aa α-helix sandwiched between β5 and β6 (Fig. 4E). Finally, the PilZNR (YcgR_2) domain of the *V. cholerae* VCA0042 (PlzD) and *B. subtilis* MotI (DgrA) proteins contains mostly the same structural elements as the PilZN domain, but its first α-helix is much shorter, only 5 amino acid residues (Fig. 4F). In summary, all these domains show clear similarity to the canonical PilZ and can be considered divergent PilZ-related domains.

### PilZ-related domains of unknown structure

The 32^nd^ release of the Pfam database (last accessed August 31^st^, 2019) listed 8,676 sequences that had PilZ as their only recognized domain (15). More than 1,800 of them consisted of 250 ± 70 amino acid residues, suggesting that, in addition to PilZ, they contained an unrecognized protein domain of a similar length. We have extracted such protein sequences from UniProt (51), clustered them by sequence similarity, and performed sequence analysis and secondary structure predictions for representatives of the largest clusters and diverse taxonomic lineages. Some of the largest clusters of proteins retrieved this way represented tPilZ domains and PilZ dimers, whose lengths also fall within the 180-320 aa range. However, this analysis also revealed several previously unrecognized domains, some of which, while not fitting the Pfam models for YcgR, YcgR_2, or DUF5634_N domains, were similar to the canonical PilZ in having the same (predicted) six-stranded β-barrel fold and likely represented additional xPilZ domains.

The list of previously and newly described non-canonical PilZ domains is presented in Table 2. Many of these domains show limited phylogenetic distribution, being found only in closely related members of a specific phylum, class, family, or just a single genus. For a lack of a better description, we have tentatively named these novel variants PilZN3 and PilZN4 (found in spirochetes and members of *Aquificae* and *Fibrobacteres*), PilZN5 (found in actinomycetes), PilZN6 (found in deltaproteobacteria), and PliZN7 (found in cellulose-degrading clostridia).

To our knowledge, *B. burgdorferi* PlzA (BB0733) and *Treponema denticola* TDE0214, which both have the PilZN3-PilZ domain architecture, are the only proteins with novel xPilZ domains that have been experimentally characterized; they both affect motility and virulence of the respective spirochetes (52-54). Secondary structure predictions indicate that the PilZN3 domain retains the C-terminal α-helix of PilZ and has three additional α-helices (Fig. S4A). While some spirochetes encode multiple PilZ-containing proteins, those with the PilZN3-PilZ architecture are the only ones found in *Borrelia*/*Borreliella* spp. and *Treponema pallidum*, the causative agent of syphilis.

*Treponema denticola* also encodes another PilZ domain protein, TDE1318, which combines a canonical PilZ domain with a 210-aa region that consists of a six-stranded xPilZ domain without the C-terminal α-helix and a unique pentahelical subdomain at the N-terminus (Fig. S4B). The first α-helix of this subdomain consists exclusively of uncharged (mostly hydrophobic) amino acid residues, indicating that it could localize to the membrane and serve as a membrane anchor for the entire protein. In addition to spirochetes, such proteins have been found in members of *Aquificae* and *Fibrobacteres*.

Finally, some new domain combinations can be found in the same genome in two or more paralogous forms. Thus, *Geodermatophilus obscurus* and several other actonobacteria encode two copies of the proteins with PilZN5-PilZ domain combination, *Desulfarculus baarsii* and other deltaproteobacteria encode two PilZN6-PilZ proteins, whereas such cellulosolytic clostridia as *Hungateiclostridium thermocellum* and *Pseudobacteroides cellulosolvens* encode, respectively, 9 and 11 proteins with the PilZN7-PilZ domain architecture (Fig. S4, panels C-E).

## DISCUSSION

In the constantly growing list of c-di-GMP-binding proteins, PilZ domain occupies a unique position, not only as the first c-di-GMP receptor to be identified and experimentally characterized (16, 19, 22-24) but, along with the recently described MshEN (11, 12) and the inactivated GGDEF and EAL domains, as the only dedicated c-di-GMP-binding domains that are truly widespread in bacteria. Other c-di-GMP receptors have a relatively narrow phylogenetic distribution; many of them likely evolved as specific add-on adaptations to c-di-GMP binding within common protein superfamilies (13, 14, 55-57).

Full-length PilZ domains consisting of a six-stranded β-barrel and a C-terminal α-helix are encoded in the genomes from numerous bacterial phyla, see Pfam entry <u>PF07238</u> or the web site https://www.ncbi.nlm.nih.gov/Complete_Genomes/c-di-GMP.html). This widespread distribution, including their presence in such early-diverging phyla as *Aquificae* and *Thermotogae*, strongly suggests that this form of PilZ is the ancestral one, which is another reason why we refer to it as the “canonical” version. Structural comparisons show the presence of essentially the same canonical variant of PilZ in BcsA, YcgR, MotI, and MrkH proteins, where PilZ is the C-terminal domain and also in Alg44, where PilZ is at the N-terminus (Fig. 1A,B). While certain stand-alone PilZ domains show disruption of the c-di-GMP-binding motifs and the resulting loss of c-di-GMP binding (Fig. 4B), PilZ domains in fusion proteins exhibit no such change and invariably retain the ability to bind c-di-GMP. It appears that the PilZ domains in BcsA, Alg44, and in YcgR- and MotI-related proteins have evolved under strict selection that ensured preservation of c-di-GMP binding and, accordingly, their participation in the c-di GMP signaling circuits.

In addition to these protein types, the PilZ domain can be found in other domain architectures, see the Pfam database (15) entry http://pfam.xfam.org/family/PilZ or Fig. 1 in refs. (22) and (10)). Although PilZ domains in those proteins have not been structurally characterized, the high degree of sequence conservation suggests that they all retain the same PilZ structure, further expanding the list of canonical PilZ domains.

Except for the two-helical insert, tPilZ, the “atypical” tetramer-forming version of the PilZ domain, is very similar to the canonical version (Fig. 2 and 4A). The tPilZ domain is found, in addition to the Proteobacteria, in members of several other phyla (Fig. S3 and Table S3). Because of the loss of c-di-GMP-binding motifs, members of the tPilZ family have often been overlooked. Thus, a comprehensive study of c-di-GMP regulation in *L. pneumophila* (58) did not even list the Lpg1926 protein as PilZ, even though this protein retains the c-di-GMP-binding motifs (Fig. S3). On the other hand, tPilZ domains of *V. cholerae* protein PlzB (VC1885) and *P. aeruginosa* PA2989 were just assumed to be the canonical PilZ domains (23, 47).

The tPilZ domain could have evolved fairly early in the bacterial evolution through an insertion into the canonical *pilZ* gene of a ∼200-bp fragment encoding helices α2 and α3. However, outside of gamma- and delta-subdivisions of *Proteobacteria* and the phylum *Nitrospirae*, tPilZ domain is found in a relatively small number of organisms with a patchy phylogenetic distribution (Table S3). Thus, this version of PilZ likely spreads by horizontal gene transfer, which implies that it carries out some beneficial function(s). As is the case with canonical PilZ, certain members of the tetrameric PilZ family retain intact c-di-GMP-binding motifs, whereas others have totally lost them (Fig. S3) and appear to participate in regulation solely through protein-protein interactions. The recognition of the tPilZ domains as a specific subfamily of PilZ should stimulate investigation of their roles in bacterial regulation. Two tPilZ proteins, *V. cholerae* PlzB and *X. campestris* Xcc6021, have been reported to contribute to virulence despite their inability to bind c-di-GMP (23, 46), making a detailed characterization of this family an interesting and promising task.

### Conservation of the C-terminal α-helix

As noted above, canonical PilZ domains differ from xPilZ ones by the presence of a long C-terminal α-helix. This helix is preserved in the PilZ fusion proteins BcsA and Alg44 and PilZ dimers, as well as in the tetrameric PilZ domain (Fig. 2). Based on sequence comparisons, it has also been predicted in PilZ-containing chemotaxis receptors Tlp1 and Aer from *Azospirillum brasilense* {Russell, 2012 #1424; O’Neal, 2019 #1796), Ser/Thr-type protein kinase Pkn1 (<u>MXAN_1467</u>) from *Myxococcus xanthus* {Munoz-Dorado, 1991 #562}, and many other PilZ fusion proteins. This helix is characterized by a high number of charged amino acid residues, particularly Arg and Lys, which makes it a prime candidate for protein-protein interactions. Indeed, in *P. aeruginosa* MapZ, this helix mediates the interaction of PilZ with the methyltransferase CheR (59). In *K. pneumoniae* MrkH, it has been implicated in binding DNA (38). In the cellulose synthase structure, the C-terminal α-helix of PilZ is split into two, which both pack against the glycosyltransferase domain (BcsA has an additional long C-terminal α-helix after the PilZ domain) (25, 60). The C-terminal α-helix also plays a role in the interaction of the PilZ domain of *X. citri* with the type IV pilus biogenesis regulator FimX (32). Finally, in the PilZ domain of Alg44, C-terminal α-helices of two monomers interact, contributing to the formation of the PilZ dimer (37). These data clearly indicate that the C-terminal α-helix of PilZ, with its clustering of charged residues, is a key element mediating protein-protein interactions by the PilZ domain.

### Role of the conserved hydrophobic residues

While the presence of conserved hydrophobic residues in the PilZ domain could be seen even in its original description (16) and is clearly visible in the HMM logo of PilZ on the Pfam database (15) web site http://pfam.xfam.org/family/PF07238#logoBlock, the role of these residues has long remained overlooked. However, strict conservation of these residues in all canonical PilZ and tPilZ domains (Fig. 2), as well as limited conservation in xPilZ domains (Fig. S2 and data not shown), are hardly accidental. These residues appear to play several important roles in maintaining the structure and activity of various PilZ-related domains. First, by forming a hydrophobic core in the middle of the protein, these residues could play a key role in the folding of the PilZ domain. Given that these residues come from five different β-strands (Fig. 2), this hydrophobic core appears to hold the entire structure together, ensuring the rigidity of the entire β-barrel. This should be important irrespective of the ability of a particular PilZ domain to bind c-di-GMP and participate in protein-protein interactions. Second, in those PilZ domains that bind c-di-GMP, the fixed position of the central *h* residue of the c-di-GMP-interacting [D/N]hSxxG motif is involved in stabilizing the SxxG β-turn hairpin, which is important for c-di-GMP binding. Conversely, c-di-GMP binding can be expected to affect the organization of the hydrophobic pocket and the position of the C-terminal α-helix, which a) provides two (sometimes three) hydrophobic residues to that pocket and b) plays a key role in protein-protein and protein-DNA interactions mediated by the PilZ domain (see above). This way, the *h* residue of the [D/N]hSxxG motif could serve as a molecular read-out, connecting c-di-GMP binding to changes in the domain interactions in at least some PilZ domains [in cellulose synthase the read-out is movement of the gating loop, while in YcgR it is likely the distance between the two PilZ domains in a dimer (22, 25, 47)]. However, verification of this proposed mechanism would require a detailed analysis of the subtle changes in high-resolution structures of PilZ with and without bound c-di-GMP.

### Pathways of PilZ evolution

The widespread distribution of the canonical c-di-GMP-binding form of PilZ and its conservation in a variety of domain contexts suggests the stand-alone form of PilZ was the ancestral version of this domain. From that, one of the pathways of PilZ evolution likely involved the insertion of a two-helical fragment that led to the formation of the tetramer-forming tPilZ domains (Fig. 5). For some reason, tetramer formation precluded domain fusions, such that tPilZ domains are almost always found in a stand-alone form. C-di-GMP binding motifs are seen in tPilZ proteins from several diverse phyla (Fig. S3, Table S3), indicating that the c-di-GMP binding form was the ancestral one. However, the loss of the c-di GMP binding motifs is a constant feature in the tPilZ family and can be traced even within a single genus, *Pseudomonas* (Fig. S3).

**Figure 5.**
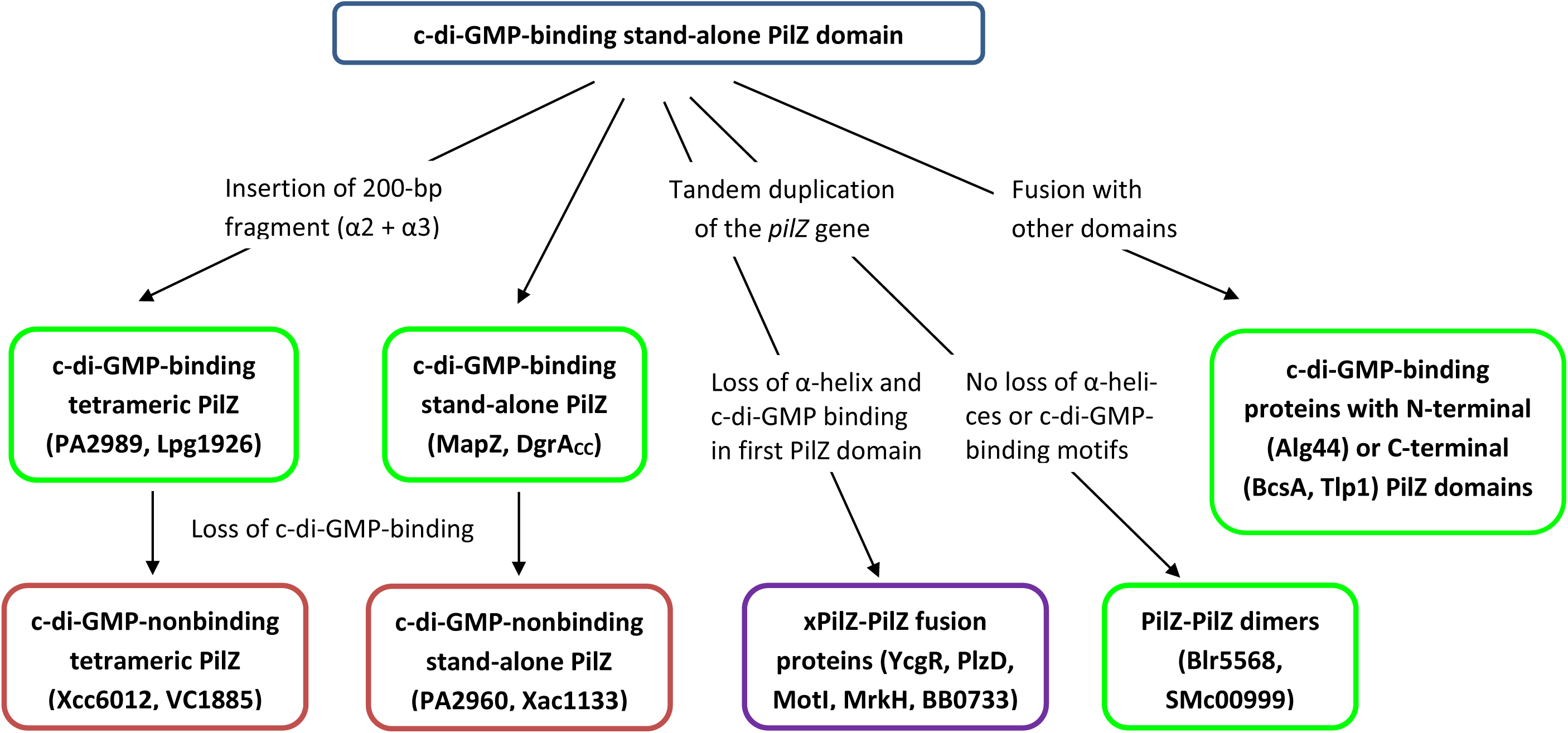
Proposed pathways of PilZ evolution. C-di-GMP-binding versions are in green frames, non-binding are in red frames, the purple frame around YcgR, PlzD, MotI, MrkH, and BB0733 indicates that only second (canonical) of the two PilZ-related domains is c-di-GMP-binding. Except for the *Caulobacter crescentus* protein DgrA_CC_ (GenBank: AAK23578, ref. 20), the proteins used as examples are described in the text or listed in Tables 1 and 2. See the text for details.

Another major pathway of PilZ evolution involved a variety of domain fusions. Some of them, such as in BcsA, Alg44, and many other proteins, preserved the canonical PilZ structure and the ability to bind c-di-GMP. A special widespread variant of such fusions was the tandem duplication of the PilZ domain, which occasionally led to the formation of fusion proteins consisting of two canonical PilZ domains, such as, for example, *B. diazoefficiens* proteins Bll4394 and Blr5568 (16) and *S. meliloti* protein SMc00999. Alternatively, tandem duplication led to the loss of the C-terminal α-helix in the N-terminal PilZ domain, often accompanied by a variety of other structural changes, and ended up in xPilZ-PilZ domain architectures. This diversification of the N-terminal xPilZ domains gave rise to PilZN, PilZNR, PilZN1, PilZN2, and other domains, some of which are listed in Table 2. All these domains have lost the c-di-GMP binding motifs, relegating c-di-GMP binding to the inter-domain linker at the beginning of the second (canonical) PilZ domain. The proposed pathways of PilZ evolution are schematically depicted in Fig. 5. We hope that this scheme will clarify the relationship between known PilZ, tPilZ, and xPilZ domains and will serve as a guide to discovering and characterizing new members of the PilZ superfamily.

## MATERIALS AND METHODS

### Structural alignment of PilZ-related domains

The available 3D structures of PilZ-related domains (Table S1) were extracted from the PDB and MMDB databases and compared using Dali, VAST and PDBeFold tools (44, 45, 61, 62). The lengths of aligned Cα traces, sequence identity in the aligned segments and their root mean standard deviation (RMSD) values were as reported by the Dali (Table 1) and VAST (Table S2) servers. Structural superpositions were generated with PyMOL (63) using the high-resolution structure of the stand-alone PilZ domain in the *P. aeruginosa* protein MapZ (PDB: 5XLY_B, residues 4-107) as the reference and aligned β-strand segments from the Dali and VAST outputs as a guide. For clarity, the second C-terminal α-helix of MapZ (residues 109-121), which is not part of the PilZ domain core, was removed from Fig. 1 and S1. Structure-based sequence alignments of various PilZ domains were taken from Dali and VAST outputs and supplemented with the alignments generated by the HHpred tool (64) of the MPI Bioinformatics Toolkit (65). The sequence logos of c-di-GMP-binding motifs were generated using the WebLogo program (66) from an alignment of 9,224 UniProt sequences that was retrieved from Pfam (15) and manually edited to remove gaps and leave only those sequences where both motifs were intact.

### Sequence analysis of tetramer-forming PilZ domains

The initial lists of tPilZ domain-containing proteins were obtained using PSI-BLAST (67) searches against the NCBI protein database and HMMer (68) searches against UniProt (51) with the *X. campestris* protein XccB100_2234 (GenBank: <u>CAP51589</u>, UniProt: <u>B0RT03</u>, PDB: <u>3RQA</u>) as the query. We further carried out database searches with PSI-BLAST and HMMer starting from the proteins in that initial list and additionally employed PHI-BLAST (67) with specified KLD and, subsequently, expanded [KR]h[DEN] motifs. The taxonomic representation of tPilZ domains was evaluated through BLAST searches against phylum-specific protein sets (excluding Proteobacteria).

### Identification of novel PilZ-related domains

The list of UniProt entries containing PilZ as their only recognized domain was taken from Pfam (15) and sorted by length to extract those sequences whose lengths fell within the 180-350 aa range. These sequences were clustered with the MMseqs2 tool (69) of the MPI Bioinformatics Toolkit (65) and representatives from the top 20 clusters were studied one-by-one. Additional clusters were included in the analyzed set based on the phylogenetic representation of their members. Selected representatives of each cluster were checked for the presence of known protein domains in their N-terminal regions using CD-Search (70) with relaxed parameters (expect value cut-off of 10) and the predicted structural similarity to PilZ was checked using HHPred (64, 65). Protein secondary structures were taken from the HHpred outputs or predicted using JPred (71).

## ACKNOWLEDGMENTS

This work was supported by the NIH Intramural Research Program at the National Library of Medicine (MYG) and by the Ministry of Education, Taiwan, ROC, under the ATU plan, and by the National Science Council, Taiwan, ROC (Grants 102-2113-M005-006-MY3 to SHC). We thank Daria Shalaeva (Osnabrück) and Yuri Wolf (Bethesda) for helpful suggestions.

## Supplementary materials

Table S1. PilZ-related domains with known 3D structure

Table S2. Structural similarity of canonical, tetrameric, and certain divergent PilZ domains

Table S3. Phylogenetic distribution of tetramer-forming PilZ domains

Figure S1. Structural superpositions of *P. aeruginosa* MapZ with canonical PilZ domains.

Figure S2. Structure-based sequence alignment of canonical PilZ domains

Figure S3. Sequence alignment of tetramer-forming PilZ domains

Figure S4. Sequence alignments of the newly identified xPilZ domains.

